# Efficient genome editing using modified Cas9 proteins in zebrafish

**DOI:** 10.1101/2023.11.12.566744

**Authors:** Laura Dorner, Benedikt Stratmann, Laura Bader, Marco Podobnik, Uwe Irion

## Abstract

The zebrafish (*Danio rerio*) is an important model organism for basic as well as applied biomedical research. One main advantage is its genetic tractability, which was greatly enhanced by the introduction of the CRISPR/Cas method a decade ago. The generation of loss-of-function alleles via the production of small insertions or deletions in the coding sequences of genes with CRISPR/Cas systems is now routinely achieved with high efficiency. The method is based on the error prone repair of precisely targeted DNA double strand breaks by non-homologous end joining (NHEJ) in the cell nucleus. However, editing the genome with base pair precision, by homology-directed repair (HDR), is by far less efficient and therefore often requires large-scale screening of potential carriers by labour intensive genotyping. Here we show that the Cas9 protein variant SpRY with relaxed PAM requirement can be used for gene targeting in zebrafish, thus expanding the versatility of the method. In addition, we demonstrate that the incorporation of an artificial nuclear localisation signal (aNLS) into the Cas9 protein variants not only enhances the efficiency of gene knock-out but also the frequency of HDR thereby facilitating the efficient modification of single base pairs in the genome. Our protocols provide a guide for a cost-effective generation of versatile and potent Cas9 protein variants and efficient gene editing in zebrafish.

## Introduction

During the last decade the CRISPR/Cas system has greatly facilitated genome editing in many organisms, including so-called ‘model organisms’, like mouse, *Drosophila* or zebrafish (Housden, Muhar et al. 2017). The system is based on an RNA-dependent endonuclease, very often the Cas9 protein from *Streptococcus pyogenes*, and an RNA molecule, the single-guide RNA (sgRNA), which provides sequence specificity through base-pairing with the target DNA (Jinek, Chylinski et al. 2012). In addition to the base pairing of RNA and DNA a short motif, the proto-spacer adjacent motif (PAM), needs to be present for enzyme activity. In the case of Cas9 from *S. pyogenes* the PAM sequence is NGG (Jinek, Chylinski et al. 2012), which leads to some constraint when searching for potential target sites in the genome of interest. An engineered variant of Cas9, containing five point mutations compared to conventional Cas9 protein and named SpRY, has a greatly reduced PAM requirement while still retaining endonuclease activity (Walton, Christie et al. 2020). The generation of knock-out alleles of protein-coding genes with the CRISPR/Cas system is most often achieved by introducing targeted DNA double-strand breaks (DSBs) in vivo and the subsequent error-prone repair by non-homologous end-joining (NHEJ), frequently leading to small deletions or insertions, which disrupt the open reading frame (Hruscha, Krawitz et al. 2013). More precise genome editing can be achieved either by employing base editors (Pickar-Oliver and Gersbach 2019) or by a mechanism called homology-directed repair (HDR), where a donor-template is provided to the cell for the repair of the DNA DSBs (Auer, Duroure et al. 2014). In eukaryotes the Cas9 protein needs to be targeted to the nucleus in order to be active. This is frequently achieved by tagging the protein with a nuclear localization signal (NLS) derived from the SV40 large T-antigen. However, it was shown that in medaka (*Oryzias latipes*) embryos the SV40 NLS does not direct efficient nuclear import during very early stages of development (Inoue, Iida et al. 2016); which is when Cas9 activity would be most effective to generate low levels of mosaicism and high chances of germ line transmission of the engineered allele. An optimized artificial NLS (aNLS) was found that leads to prominent nuclear localization of GFP immediately after fertilization in medaka (Inoue, Iida et al. 2016). In zebrafish (*Danio rerio*) the inclusion of this aNLS sequence into a bi-partite hei-tag also boosts genome editing efficiency of Cas9 protein and of cytosine-to-thymine base editors (Thumberger, Tavhelidse-Suck et al. 2022). Here, we show that the SpRY variant of Cas9 can be used for efficient genome editing in zebrafish. However, we find that not all sites we tested are targeted with the same efficiency and that individual target sequences might be of great importance and probably need to be assessed on a one-by-one basis. In addition, we show that the addition of an aNLS to Cas9 protein enhances its efficiency to a degree that allows the precise exchange of one codon in the genome of zebrafish without the need for a visible read-out or pre-selection of the F_0_ larvae. We use the method to precisely change the coding sequence for *kcnj13* in zebrafish into the sequence found in *Danio aesculapii*, its closest sister species (Podobnik, Frohnhöfer et al. 2020, Podobnik, Singh et al. 2023). Zebrafish carrying the edited sequence do not show any visible phenotype, thus demonstrating that the Kcnj13 proteins from both species are functionally equivalent. We describe a very simple and robust method that makes use of purified proteins and in vitro transcribed RNAs, and thus has the potential to be used in many laboratories for genome editing, and even for teaching undergraduate students in courses.

## Results

### Efficient *albino* knock-out with the SpRY protein

To assess whether a modified version of the Cas9 protein (SpRY), which has been reported to allow near PAM-less genome targeting in cultured human cells (Walton, Christie et al. 2020), is also active in zebrafish we used the *albino* (*alb/slc45a2*) gene as target. Mutations in *alb* can be generated with very high efficiency using the conventional Cas9 protein, and it is possible to repair a premature stop codon in the *alb*^*b4*^ mutant by co-injection of a donor-plasmid for homology-directed repair (Irion, Krauss et al. 2014). We generated six sgRNAs to target different positions in three exons of *alb* (Fig.1 A) and injected them together with a purified fusion protein of SpRY with C-terminal mCherry, N-terminal Strep- and His-tags for purification and an SV40 large T NLS (for simplicity referred to a SpRY in the rest of the manuscript), into TU embryos. For the evaluations of these *alb* k.o. experiments we examined the mosaic F_0_ larvae after 3 days and classified them into five categories according to the severity of the *alb* phenotype (Fig.1 B-F’): 1 (none), no loss of pigmentation discernible; 2 (weak), loss of pigmentation in individual cells, usually only clearly visible in the retinal pigment epithelium; 3 (moderate), regional loss of pigmentation clearly visible; 4 (good), maximum of 50 pigmented cells left; 5 (very good/complete), maximum of five pigmented cells. For three of the tested target sites (U1, U3, U5) we found no or only very weak activity of the injected SpRY-RNP. In two cases (U2 and U4) we found moderate activity, but in one case (OP2) the activity was high, with 49% to 82% (61.6% average) of the larvae falling into categories 4 and 5 (Fig.1 G, Table 1). This is comparable to the k.o. activity we regularly achieve with unmodified Cas9 protein.

**Figure 1:**
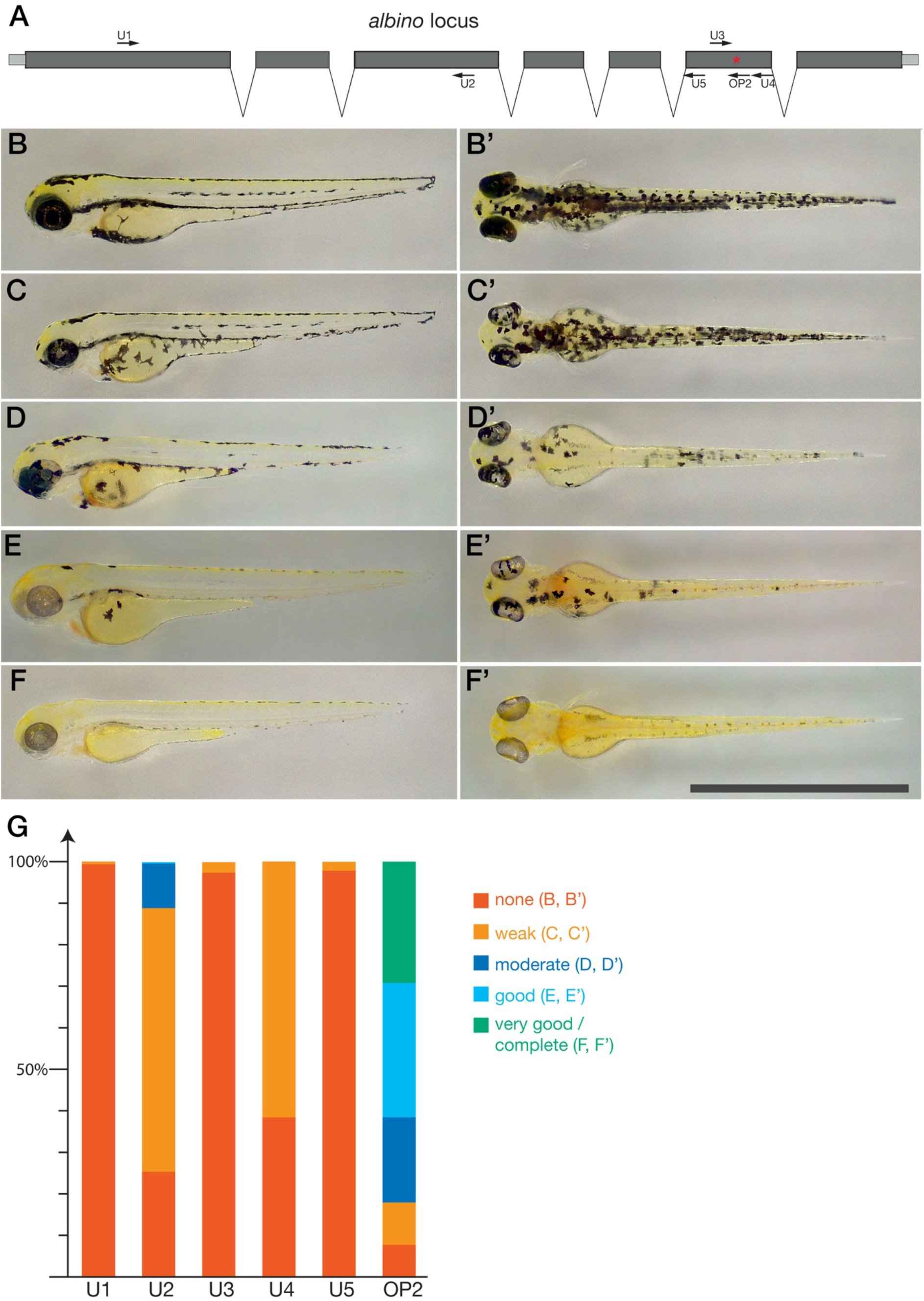
Efficient knock out of the *albino* gene using the SpRY protein. In (A) the *albino* locus is schematically shown, exons are in grey, introns are shown as gaps, not to scale. The coding sequence is in dark grey and the different target sites in exons 1, 3 and 6 are indicated. The different categories for the evaluation of the knock-out efficiency are shown in (B) - (F’). Lateral views (B) - (F) and dorsal views (B’) - (F’) of larvae 3 dpf are shown. Category 1, no knock-out (none) (B), (B’), category 2 weak (C), (C’), category 3 moderate (D), (D’), category 4 good (E), (E’) and category 5 very good/complete knock-out (F), (F’). In (G) the results for the six target sites tested are shown. For detailed results see Table 1.

**Table 1:**
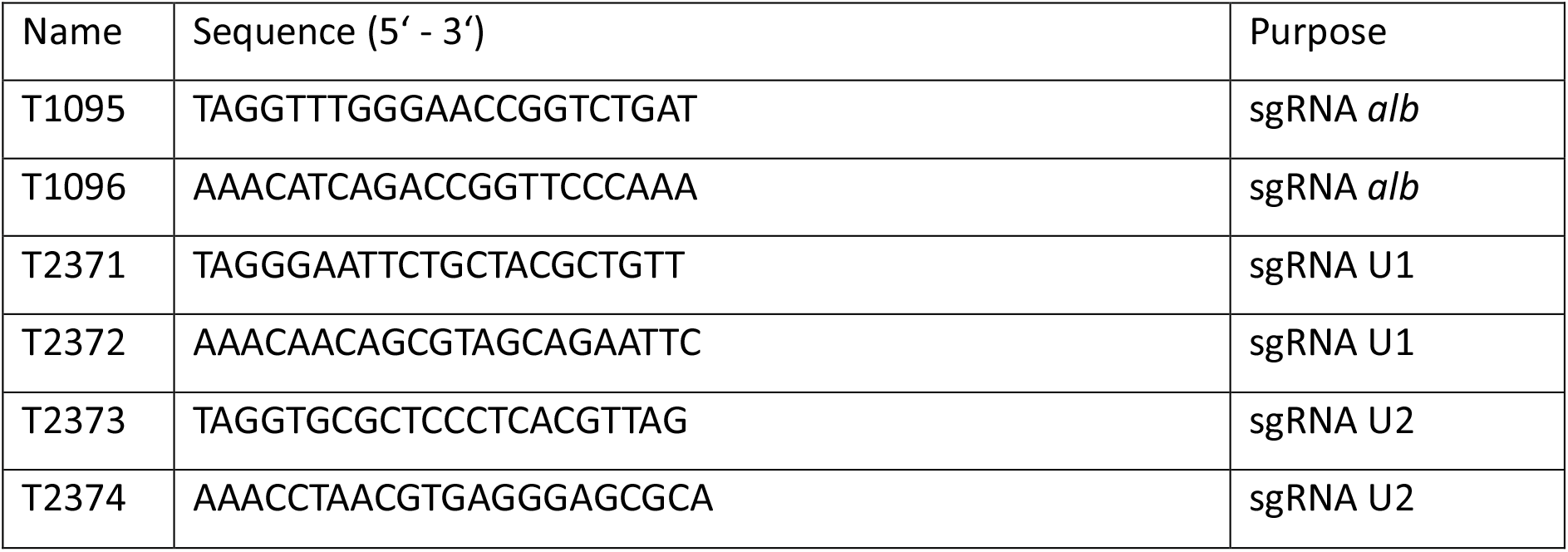
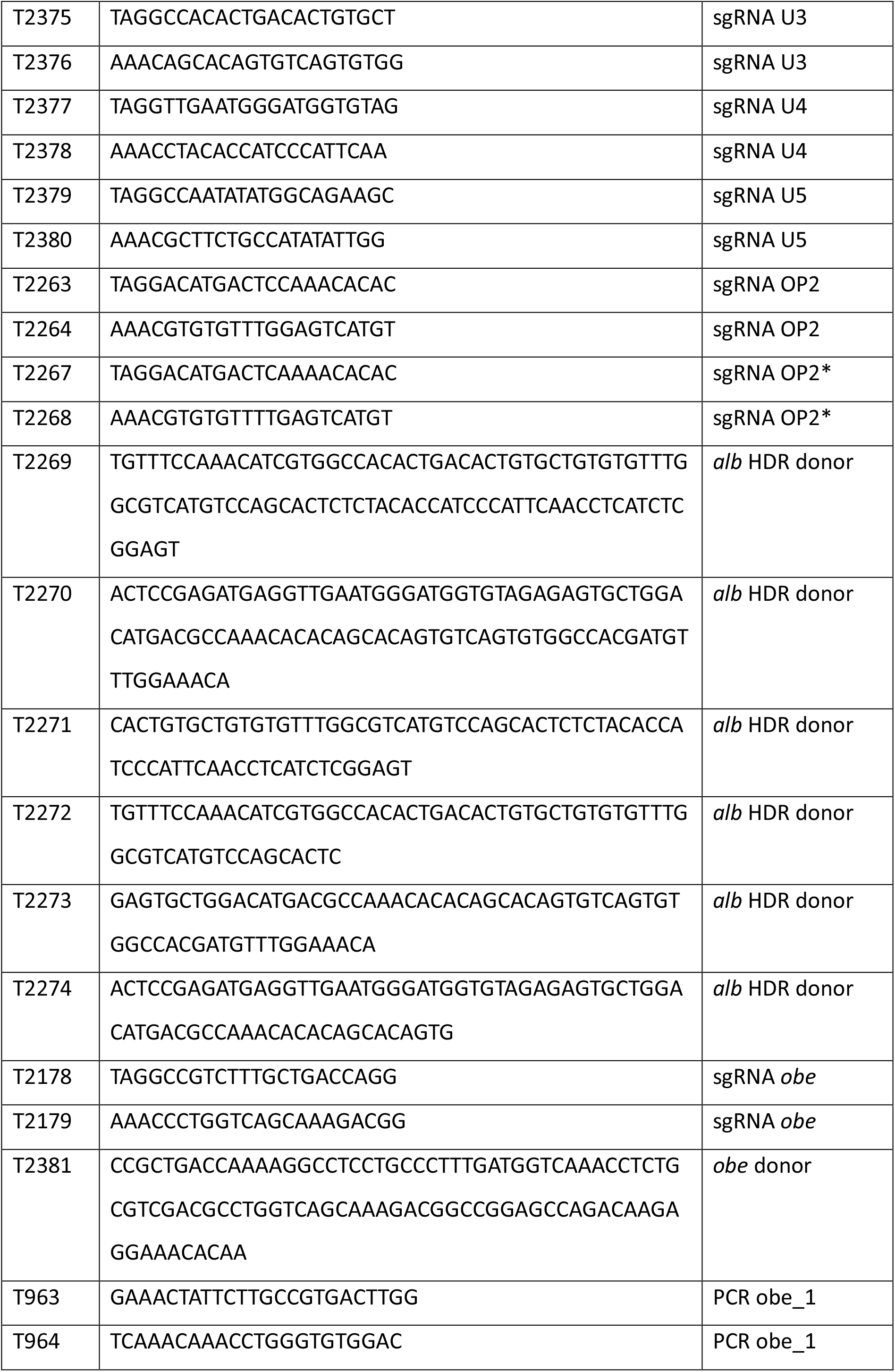
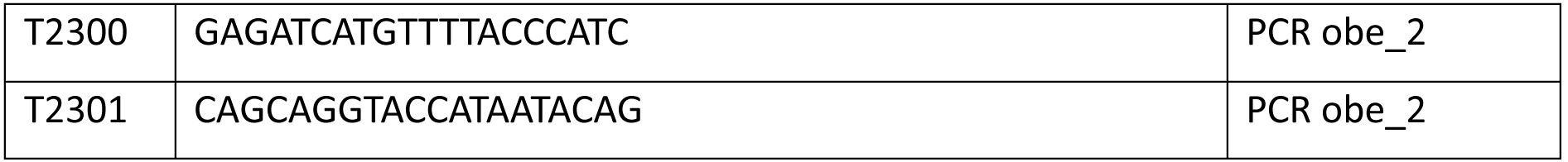
Oligonucleotides.

### Repair of the *albino*^*b4*^ mutation with the SpRY protein and oligonucleotide donors

As the target site for the OP2 sgRNA overlaps with the mutation in *alb*^*b4*^, a premature stop codon in exon 6 (Fig.2 A) (Dooley, Schwarz et al. 2013), we decided to co-inject SpRY and the OP2* sgRNA with oligonucleotides as donor templates for homology-directed repair of the mutation (Fig.2 B). To evaluate these repair experiments we grouped the larvae into four categories: 1 (none), no pigmented cells visible; 2 (weak), fewer than 15 pigmented cells; 3 (moderate), 15 to 25 pigmented cells; 4 (good), more than 25 pigmented cells (Fig.2 C-F’).

**Figure 2:**
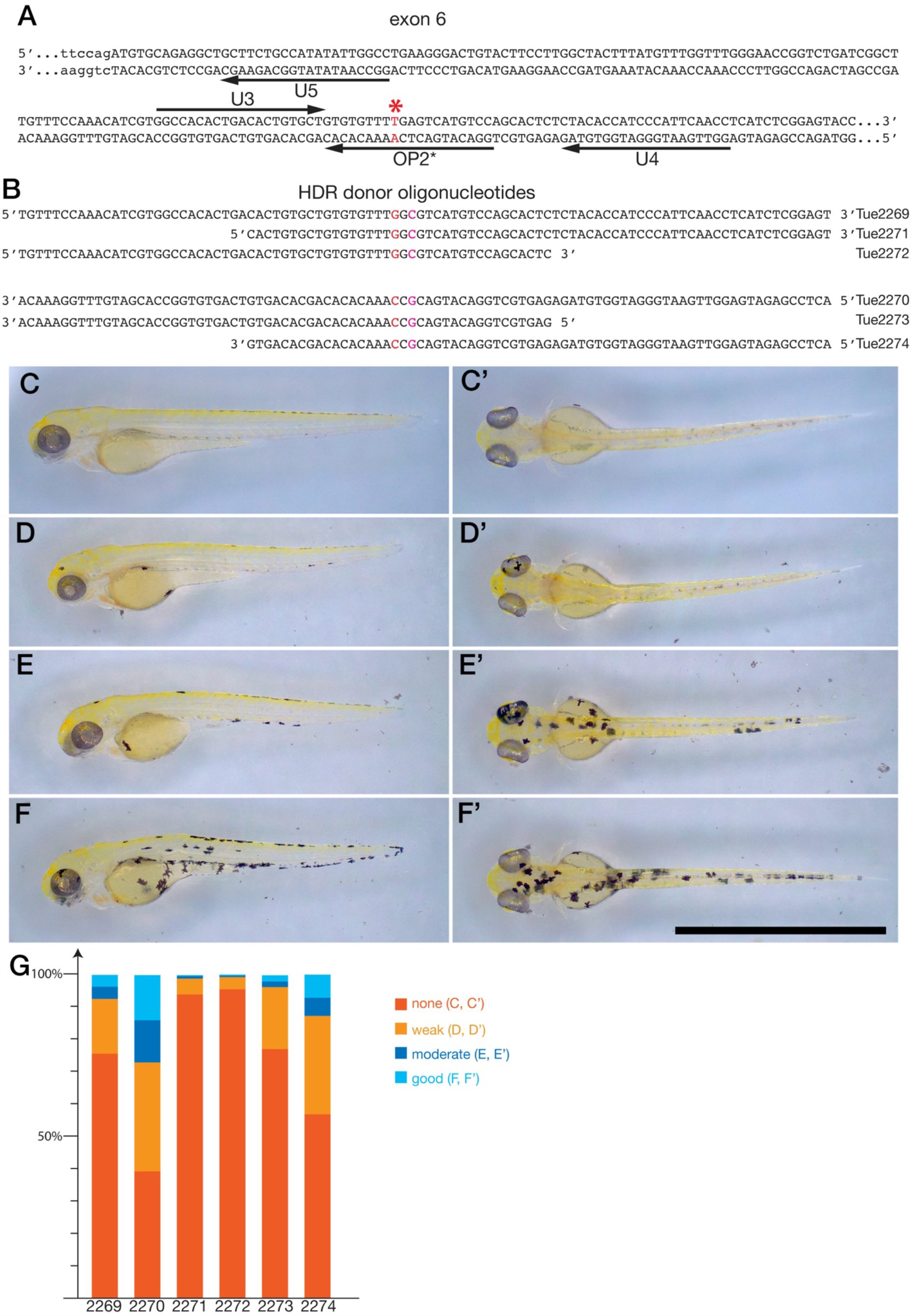
Homology directed repair of the *albino*^*b4*^ mutation using the Cas9 variant SpRY. In (A) the partial sequence of exon 6 of the *albino* gene is depicted, coding sequence in capital letters, intron sequence in small letters. The target sites U3, U4, U5 and OP2* are indicated by arrows; the mutation in *alb*^*b4*^ leading to a premature stop codon is shown in red and marked with an asterisk. In (B) the oligonucleotides used as donors for HDR are shown. The two base pairs that are altered are shown in red; note the 5’ and 3’ ends of the oligonucleotides indicating the DNA strand they correspond to. The different categories for the evaluation of the repair efficiency are shown in (C) - (F’). Lateral views (C) - (F) and dorsal views (C’) - (F’) of larvae 3 dpf are shown. Category 1, no repair (none) (C), (C’), category 2 weak (D), (D’), category 3 moderate (E), (E’) and category 4 good repair (F), (F’). In (G) the results for the six tested donor oligonucleotides are shown. For detailed results see Table 2.

Using oligonucleotides of different lengths, we found that one of them (TÜ2270) leads to very high numbers (up to 66%) of larvae with pigmented cells, indicative of the repair of the *alb*^*b4*^ mutation (Fig.2 G). This donor oligonucleotide corresponds to the DNA sense-strand with regard to the sgRNA molecule. Other oligonucleotides, either shorter in length or corresponding to the antisense DNA strand, lead to lower HDR efficiencies (Fig. 2 G, Table 2). However, most of the larvae we obtained in this experiment show only very few pigmented cells and thus fell into categories 2 and 3, much fewer (max. 16%) larvae fell into category 4 with more than 25 pigmented cells (Fig.2 G). When we raised these fish of category 4 to adulthood and crossed them to *alb*^*b4*^ mutants we found that only one fish out of 16 showed germ-line transmission of the repaired allele.

### Enhanced efficiency by addition of an aNLS

We hypothesized that the low efficiency of HDR, despite the relatively high frequency (up to 66% of F_0_ larvae with some pigmented cells), might be due to inefficient nuclear import of the Cas9 protein during early stages of embryonic development. For medaka it was shown that an aNLS (sequence: PPPKRPRLD) improves nuclear import of proteins during early stages of development compared to the most commonly used SV40-NLS (sequence: PKKKRKV) (Inoue, Iida et al. 2016). In zebrafish the inclusion of the aNLS into a hei-tag increased the efficiency of different Cas9 protein variants with a reduction in allele variance, indicating activity during early developmental stages (Thumberger, Tavhelidse-Suck et al. 2022). We added the aNLS sequence to the N- and C-termini of mCherry tagged Cas9 and SpRY proteins and compared the activity of the resulting fusion proteins (aNLS-Cas9 and aNLS-SpRY) with the ‘normal’ Cas9 or SpRY, which both carry an SV40NLS C-terminally. We found that addition of the aNLS sequence leads to a very obvious increase in the knock-out efficiency for the Cas9 protein with more than 85% of the injected F_0_ larvae showing no melanophore pigmentation (Fig.3 A, Table 3). However, in the case of the SpRY protein the addition of the aNLS did not result in an improvement of the k.o. efficiency (Fig.3 A). Nevertheless, using the aNLS-SpRY protein resulted in a substantial increase of the efficiency for HDR-mediated repair of the *alb*^*b4*^ mutation. We found that the number of larvae in category 4, with more than 25 pigmented cells, increased from 9.1% to 19.8% (Fig.3 C, Table 4).

**Figure 3:**
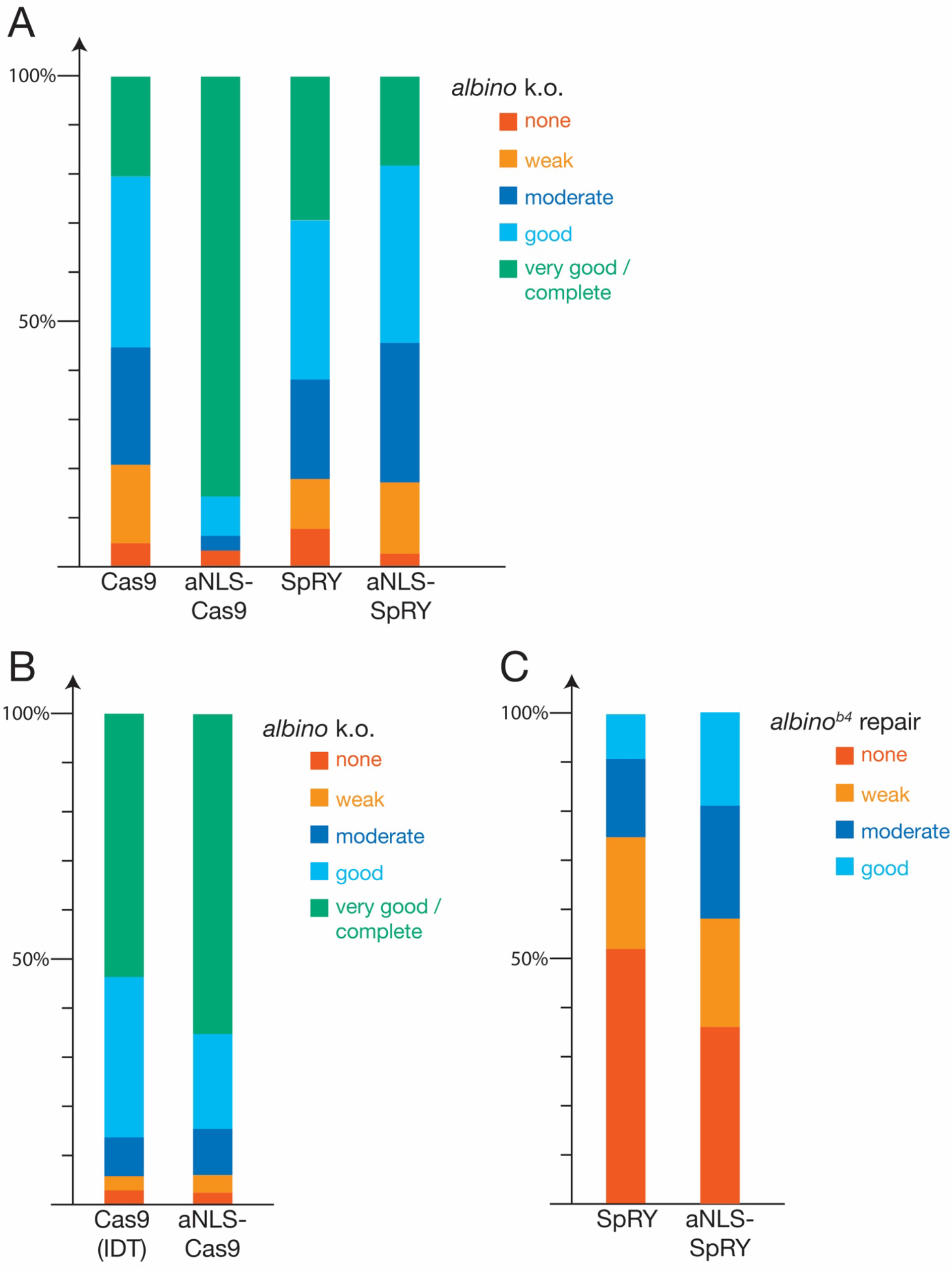
Improvement of knock-out and repair efficiencies by adding an aNLS. In (A) the efficiencies of knocking out the function of the *albino* gene are compared between Cas9 and aNLS-Cas9 and SpRY and aNLS-SpRY. Whereas in the case of Cas9 the addition of an aNLS leads to a significantly higher efficiency, 93.5% in the best two categories compared to 55.1%, this is not the case for SpRY. Detailed results see Table 3. Purified aNLS-Cas9 performs as well as commercially available Cas9 protein (Alt-R™ S.p. Cas9 V3, IDT) when tested for the knock-out of the *albino* gene (B). In addition, the repair efficiency for aNLS-SpRY is significantly higher compared to SpRY, with 19% v 9.1% in the category ‘good’ (C). For detailed results see Table 4.

### Comparison with commercial Cas9 protein

To compare our own purified aNLS-Cas9 protein with a commercially available protein (Alt-R™ S.p. Cas9 V3, IDT) we injected both together with *alb* sgRNA into wt embryos. We found that when using 500 ng/μl of the protein, k.o. mutations in *alb* were induced with comparable very high frequencies in both cases, with more than 50% of the larvae falling into category 5 (Fig.3 B, Table 5). At concentrations of 1000 ng/μl mortality rates for our own protein were too high to allow meaningful comparisons.

### Allele exchange by HDR at the *kcnj13* locus

Finally, we employed the HDR method for the directed exchange of one specific base pair in the genome of zebrafish. We targeted the first coding exon of *obelix* (*obe*/*kcnj13*) and co-injected an oligonucleotide to change one codon, CAG (Gln^23^), to CTG (Leu^23^), which is the sequence found in *Danio aesculapii*, the closest sister species to *Danio rerio* (Podobnik, Frohnhöfer et al. 2020, Podobnik, Singh et al. 2023). As we have no visible read out in this case we designed the donor-oligonucleotide to contain a second base change that will not affect the coding potential but introduce a novel target site for the restriction enzyme SalI (Fig.4 A). We screened some of the injected F_0_ larvae by PCR and restriction enzyme digest to make sure that the desired HDR outcome was present. While Cas9 protein without aNLS did not yield any positive results, in the case of aNLS-Cas9 we could detect the presence of a SalI restriction site in six PCR-fragments from 16 larvae tested. Sequencing of the cloned PCR-products confirmed that the allele exchange had taken place. However, we found that other changes near the target site had also occurred rather frequently and just one out of four cloned DNA fragments had only the desired changes. We raised approx. 100 of the injected F_0_ larvae to adulthood and then tested sperm from 64 adult male fish for the presence of the engineered allele in the germ cells. This led to the identification of eight candidate fish. When outcrossed to wild type females we identified the presence of the SalI site in F_1_ offspring of six of these candidate fish. Sequencing showed that in one case the intended DNA changes were present with no further mutations detectable, whereas in the other five cases additional mutations had been introduced. The heterozygous F1 fish, which carry the engineered allele with only the intended changes, are indistinguishable from their wild type siblings (Fig.4 C,D). In addition, we crossed the identified founder fish to a *kcnj13* mutant female and identified trans-heterozygous individuals in the resulting offspring. These fish, carrying a *kcnj13* loss-of-function allele and the engineered *kcnj13* allele, also show a wild type phenotype and are indistinguishable from heterozygous siblings (Fig.4 E,F). These results show that the proteins from both species can perform the same functions in zebrafish.

**Figure 4:**
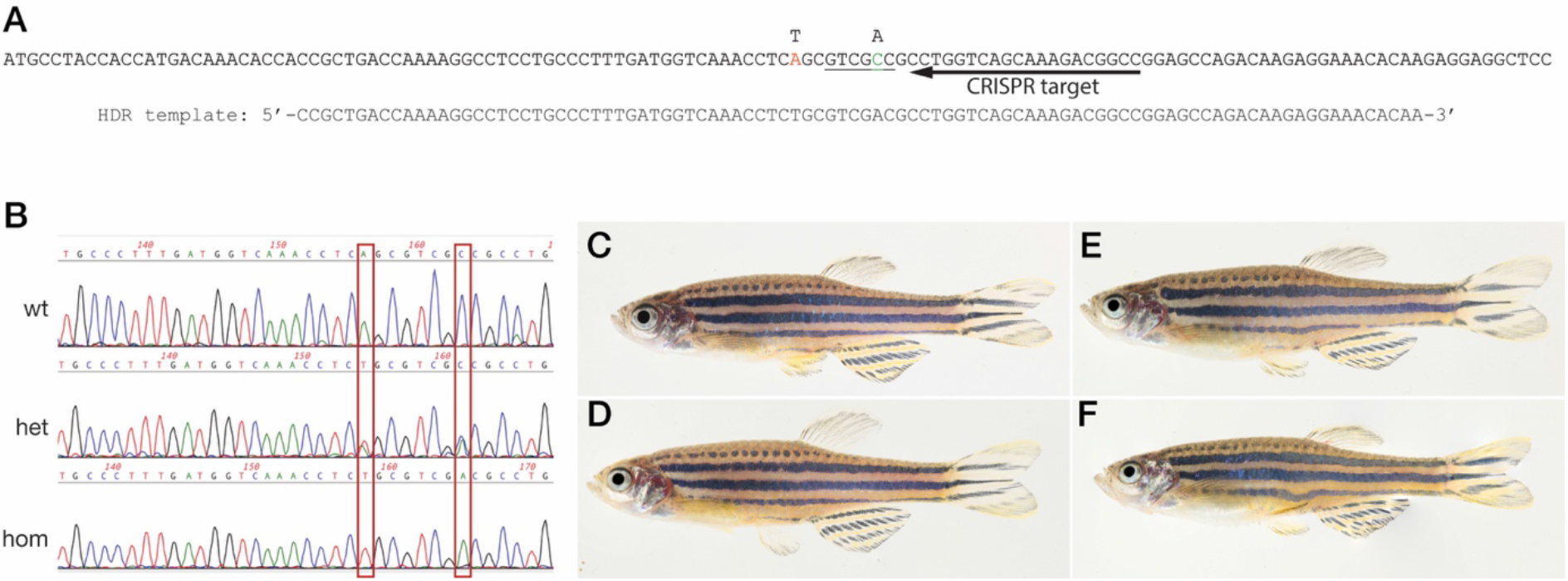
Allele exchange in the *obelix* gene. In (A) the start of the coding sequence for *obe* is shown. The CRISPR target site is indicated by an arrow, the two point mutations introduced by HDR are shown in red and green, respectively, the recognition site for SalI that is generated by the HDR is underlined. The oligonucleotide used as donor DNA is also depicted. In (B) sequences from wild type fish, and fish heterozygous or homozygous for the engineered allele are shown, the two positions that were engineered are boxed. F1 fish are shown in panels (C) to (F). There are no discernible differences between wild type siblings (C) and fish heterozygous for the engineered allele (D), nor between fish that carry a *kcnj13* loss-of-function allele over a wild type allele (E) or over the engineered allele (F).

## Discussion

Genome engineering with the CRISPR/Cas9 system in zebrafish is very robust and can be used by any researcher with only minimal prior training (Uribe-Salazar, Kaya et al. 2022). This makes the method ideal for teaching purposes, especially when combined with an assay that allows visible quantification of k.o. efficiencies after only two or three days, as is the case for the *alb* gene. While we found some variability in our results (see for example Table 3), we could reproduce all of them in several independent experiments. The observed variability most likely depends on a combination of different factors, such as the purity of the CRISPR components, mainly the different sgRNAs and oligonucleotides used, which were often different between experiments, whereas the proteins were purified once and stored as frozen aliquots. In addition, with more training the experimenters generally get better at performing injections and a higher proportion of the injected embryos survive. Our purified aNLS-Cas9 protein performed marginally better than a commercially available Cas9 (Alt-R™ S.p. Cas9 V3, IDT) when injected at our standard protein concentration of 500 ng/μl, which corresponds to 3.1 μM for Alt-R™ Cas9, a slightly lower concentration than the 4 μM recommended by the company. At the higher concentration of 1000 ng/μl, mortality rates were very high for the embryos injected with our own purified aNLS-Cas9 making a meaningful comparison impossible. A further purification step, for example by gel filtration, might improve the purity of the protein and lead to better results.

A number of different CRISPR-based methods exist for precise base pair changes in eukaryotic genomes (Anzalone, Koblan et al. 2020). Besides HDR, two alternative methods are prime editing (Anzalone, Randolph et al. 2019, Porto, Komor et al. 2020). While prime editing allows the introduction of a variety of different genomic changes, it requires the careful design and synthesis of relatively long pegRNAs and the selection of the appropriate prime editor protein (Doman, Pandey et al. 2023), which makes the method technically more challenging than HDR. Base editors, on the other hand, are active in just a small editing window, limiting potential target sites. A constraint that might be overcome by the use of near PAM-less editors (Rosello, Serafini et al. 2022) or the *de novo* generation of the desired PAM in the genome (Pakari, Wittbrodt et al. 2023), which is again more laborious than genome editing via HDR. But, most importantly, base editors only produce certain base changes, either C to T for cytosine base editors, or A to G for adenine base editors. This would not allow the changes we wanted to introduce into the *alb* or *kcnj13* genes.

Therefore, HDR is still a very convenient method for precise base changes in the zebrafish genome, especially when synthetic oligonucleotides can be co.injected as donor templates (Carrington, Ramanagoudr-Bhojappa et al. 2022). It is, however, desirable to find a target site for the endonuclease as close as possible to the position that should be changed. To test if Cas9 variants with reduced PAM-specificity can be used in zebrafish as a possibility to expand the target range of the CRISPR system, we tested SpRY. This protein is a variant of Cas9 that was engineered to have a less stringent requirement for the PAM sequence and was found to be nearly PAM-less (Walton, Christie et al. 2020). We found that for most of the target sites we tested in the *alb* gene the activity of SpRY was very low or undetectable. However, in one case we could detect a k.o. efficiency that was comparable to the high efficiency we regularly obtain using normal Cas9 protein (Irion, Krauss et al. 2014). While we did not systematically test if these differences in efficiency depend on the different PAMs, we realise that two of our target sites, OP2 and U5, share the same PAM sequence, NGC, but are targeted by SpRY with very different efficiencies (see Fig.1 G). This argues for additional constraints on the activity of the protein sgRNA complex. Our results are in good agreement with published data showing that the efficiency of SpRY seems to be target site dependent in zebrafish (Liang, Zhang et al. 2022). In conclusion, we find that SpRY might be a good option to expand the targeting range of the CRISPR/Cas system in zebrafish, but potential target sites need to be evaluated individually.

In our initial experiments using SpRY to repair the premature stop codon in the *alb*^*b4*^ mutant via HDR with oligonucleotides as donor DNA we found that the majority of all larvae only had very few pigmented melanophores. This indicates that the repair happens only after a lag phase of probably several hours after injection, otherwise we would expect larger clones of pigmented cells with the edited DNA. The reason for this might be the inefficient transport of the RNP into the nucleus during very early zygotic development, as has been shown to be the case in medaka for proteins carrying the SV40-NLS (Inoue, Iida et al. 2016). Indeed, the addition of an artificial nuclear localization signal (aNLS) to the Cas9 protein markedly increased the number of injected larvae that showed a complete *alb* phenotype to over 85% in some of our experiments, with the average being over 65%. Similarly, the repair efficiency was also higher when using aNLS-SpRY (19.0%) compared to normal SpRY (9.1%). This marked increase in efficiency prompted us to use the system for gene editing a position in the zebrafish *kcnj13* gene where a directly visible phenotypic read-out for the edit in larvae does not exist. We employed the method to produce an allele of *kcnj13* in zebrafish, which carries a point mutation that leads to an exchange of Gln^23^ to Leu, thus making it identical to the wild type *kcnj13* allele from *Danio aesculapii*. While *kcnj13* is involved in the generation of the distinct pigmentation patterns in both species, it was shown that the gene function has diverged between them (Podobnik, Frohnhöfer et al. 2020). We find that in zebrafish both proteins function identically, which demonstrates that the divergence cannot be due to changes in the coding sequence, but must be based on cis-regulatory evolution, as has been strongly suggested previously (Podobnik, Singh et al. 2023).

In summary, we show that the versatility of the CRISPR-system can be extended by the use of the SpRY protein variant and that the inclusion of an artificial NLS enhances the efficiency for the production of knock-outs as well as knock-ins via HDR.

## Materials and Methods

### Fish husbandry and micro-injections

Zebrafish were bred and maintained as described (Brand, Granato et al. 2002). *D. rerio* wild type Tuebingen (TU), *albino*^*b4*^ (Chakrabar., Streisinger et al. 1983), and *kcnj13*^*t24uI*^ (Podobnik, Frohnhöfer et al. 2020) strains were used.

The sgRNAs for the experiments were produced as described previously (Irion, Krauss et al. 2014); for the oligonucleotides used see Table 1. The standard mix for micro-injections contained 500 ng/μl Cas9 or SpRY protein, 35 ng/μl in vitro transcribed sgRNA (MegaScript T7 kit, Invitrogen), 30 ng/μl single-stranded DNA oligonucleotides (for repair experiments) and 0.05% Phenol Red in PBS + 300 mM NaCl, 150 mM KCl. One-cell stage embryos were injected with approx. 2 nl of this mix using a pneumatic system (pneumatic pico pump, WPI) and glass capillaries (Zebrafish-injection-pipettes blunt, BioMedical Instruments). Scoring of the *albino* phenotype was done at 3 days-post-fertilization (dpf).

### PCR and genotyping

For genotyping, either individual larvae at 1 dpf, still inside the chorion, or sperm, extracted from anaesthetized males by applying gentle pressure and collected with a glass capillary, were put in 100 μl 50 mM NaOH. After incubation at 95°C for 30 min 10 μl of Tris-HCl pH 8 was added, and 1 μl was used as template for the subsequent PCR amplification with the DreamTaq Hot Start DNA Polymerase (Thermo Fisher scientific). To achieve reliable amplification of the region of interest from the *obe* coding sequence a nested PCR was done. In the first round primers T963 and T964 were used, in a second round 1 μl of the first PCR was used as template with primers T2300 and T2301. For both rounds the following PCR conditions were used.

Step 1: 95°C for 2 min

Step 2: 95°C for 20 sec

Step 3: 62°C* for 20 sec

Step 4: 72°C for 30 sec

go to step 2 for 10 x

Step 5: 95°C for 20 sec

Step 6: 58°C for 20 sec

Step 7: 72°C for 30 sec

go to step 5 for 35 x

Step 8: 72°C for 3 min

Step 9: 8°C forever

*: -0.4°C per cycle

The PCR products were analysed by agarose gel electrophoresis, restriction digest with FastDigest SalI (Thermo Scientific) and Sanger sequencing (Genewiz, Azenta Life sciences).

### Protein expression and purification

The different Cas9 proteins were expressed as fusions with C-terminally attached fluorescent proteins (mCherry or mVenus) in *E*.*coli* BL21(DE3)pLysS and purified using a double tag, 6xHis and twin-StrepTag. Briefly: the bacteria were transformed with one of the plasmids, NLS-Cas9-mCherry (GenBank: OP243709), NLS-SpRY-Venus (GenBank: XXX), aNLS-Cas9-mCherry (GenBank: XXX), or aNLS-SpRY (GenBank: XXX), and grown overnight at 37°C on LB plates containing 50 μg/ml kanamycin. The bacterial colonies were scraped of the plates the next day, resuspended in 50 ml of LB medium, and used to inoculate 400 ml LB+kanamycin (starting OD_600_ of 0.1 – 0.2). The cultures were grown at 37°C with vigorous shaking until the OD600 reached 0.6 and then transferred to 18°C for 1 h. Protein expression was induced by the addition of IPTG (final concentration 0.5 mM) and the bacteria were grown for 18 h at 18°C and then harvested by centrifugation. The bacterial pellet was resuspended in 50 ml lysis buffer (20 mM Na-phosphate, 20 mM Tris-HCl, 600 mM NaCl, 150 mM KCl, 0.5 mM TCEP, 0.1% Tween20, pH=8) and the bacteria were gently lysed by freezing and thawing five times, then adding lysozyme to a final concentration of approx. 0.2 mg/ml and incubation on ice for 1 h. After centrifugation (4°C for 1 h at 3220 g) the supernatant was filtered (1.2 μm filter) and then incubated for 2 h on ice on a rocking platform with 1 ml Protino Ni-NTA Agarose (MN, Macherey-Nagel GmbH, Düren, Germany) pre-equilibrated with lysis buffer. The agarose with the bound protein was recovered by centrifugation (4°C for 5 min at 900 g), washed twice with 20 ml wash buffer 1 (= lysis buffer + 20 mM imidazole) and then at room temperature packed into a 10 ml Polyprep Chromatography Column (Bio-Rad Laboratories, Feldkirchen, Germany). The protein was eluted from the column with 5 ml elution buffer 1 (= lysis buffer + 500 mM imidazole). This eluate was applied on a second column, 1 ml StrepTactin (IBA Lifesciences, Göttingen, Germany) pre-equilibrated with lysis buffer, by gravity flow. The column was washed with 20 ml wash buffer 2 (= IBA Buffer W + 300 mM NaCl) and the protein eluted with 1 ml elution buffer 2 (= IBA Buffer E + 300 mM NaCl). The eluted protein was the dialyzed (SnakeSkin Dialysis Tubing, MWCO 10,000, Thermo Scientific) overnight using 3 x 2 l of PBS + 300 mM NaCl and 150 mM KCl. The protein concentration was adjusted to 1 (or 2) mg/ml and 10 μl aliquots were frozen at - 70°C.

For comparisons Cas9 protein (Alt-R™ S.p. Cas9 V3, glycerol-free, 10 mg/ml (62μM)) was purchased from Integrated DNA Technologies, Leuven, Belgium.

## Acknowledgements

We thank Roberta Occhinegro for excellent technical support and Christiane Nüsslein-Volhard for invaluable discussions and comments.

## Author contributions

Conceptualization and design of the experiments was done by UI. All authors performed the experiments and analysed the data. UI wrote the original draft of the manuscript and all authors were involved in reviewing and editing. UI acquired funding.

## Supplementary Tables

**Table S1:** *albino* k.o.; SpRY protein, different sgRNAs

|  | none | weak | moderate | good | very good /<br>complete |
| --- | --- | --- | --- | --- | --- |
| **U1** | 165 | 5 | 0 | 0 | 0 |
|  | 174 | 0 | 0 | 0 | 0 |
|  | 163 | 0 | 0 | 0 | 0 |
|  | 159 | 1 | 0 | 0 | 0 |
|  | 3 | 0 | 0 | 0 | 0 |
|  | 106 | 0 | 0 | 0 | 0 |
|  | 243 | 0 | 0 | 0 | 0 |
| **sum** | 1013 | 6 | 0 | 0 | 0 |
| % | 99.4 | 0.6 | 0 | 0 | 0 |
| N=1019 |  |  |  |  |  |
| **U2** | 17 | 33 | 2 | 0 | 0 |
|  | 20 | 32 | 2 | 0 | 0 |
|  | 17 | 113 | 3 | 0 | 0 |
|  | 24 | 108 | 11 | 0 | 0 |
|  | 13 | 25 | 2 | 0 | 0 |
|  | 29 | 27 | 3 | 0 | 0 |
|  | 44 | 74 | 48 | 2 | 0 |
| **sum** | 164 | 412 | 71 | 2 | 0 |
| % | 25.3 | 63.5 | 10.9 | 0.3 | 0 |
| N=649 |  |  |  |  |  |
| **U3** | 197 | 19 | 0 | 0 | 0 |
|  | 162 | 0 | 0 | 0 | 0 |
|  | 157 | 0 | 0 | 0 | 0 |
|  | 153 | 2 | 0 | 0 | 0 |
|  | 62 | 0 | 0 | 0 | 0 |
|  | 0 | 0 | 0 | 0 | 0 |
|  | 75 | 0 | 0 | 0 | 0 |
| **sum** | 806 | 21 | 0 | 0 | 0 |
| % | 97.5 | 2.5 | 0 | 0 | 0 |
| N=827 |  |  |  |  |  |
| **U4** | 49 | 188 | 0 | 0 | 0 |
|  | 102 | 134 | 0 | 0 | 0 |
|  | 31 | 109 | 0 | 0 | 0 |
|  | 42 | 101 | 0 | 0 | 0 |
|  | 27 | 0 | 0 | 0 | 0 |
|  | 23 | 13 | 0 | 0 | 0 |
|  | 71 | 8 | 0 | 0 | 0 |
| **sum** | 345 | 553 | 0 | 0 | 0 |
| **%** | 38.4 | 61.6 | 0 | 0 | 0 |
| N=898 |  |  |  |  |  |
| **U5** | 137 | 13 | 0 | 0 | 0 |
|  | 152 | 0 | 0 | 0 | 0 |
|  | 108 | 0 | 0 | 0 | 0 |
|  | 107 | 1 | 0 | 0 | 0 |
|  | 46 | 0 | 0 | 0 | 0 |
|  | 36 | 0 | 0 | 0 | 0 |
|  | 68 | 0 | 0 | 0 | 0 |
| **sum** | 654 | 14 | 0 | 0 | 0 |
| **%** | 97.9 | 2.1 | 0 | 0 | 0 |
| N=668 |  |  |  |  |  |
| **OP2** | 0 | 1 | 14 | 20 | 20 |
|  | 0 | 0 | 19 | 16 | 18 |
|  | 1 | 5 | 4 | 16 | 2 |
|  | 5 | 8 | 15 | 58 | 77 |
|  | 43 | 51 | 79 | 97 | 69 |
| **sum** | 49 | 65 | 131 | 207 | 186 |
| **%** | 7.7 | 10.2 | 20.5 | 32.4 | 29.2 |
| N=638 |  |  |  |  |  |

**Table S2:** *albino* repair; SpRY protein, OP2* sgRNA, different donor oligonucleotides

|  | none | weak | moderate | good | very good /<br>complete |
| --- | --- | --- | --- | --- | --- |
| **TÜ2269** | 65 | 15 | 3 | 8 |  |
|  | 69 | 5 | 7 | 3 |  |
|  | 45 | 15 | 2 |  |  |
|  | 98 | 27 | 2 | 2 |  |
| sum | 277 | 62 | 14 | 13 |  |
| % | 75.7 | 16.9 | 3.8 | 3.6 |  |
| N=366 |  |  |  |  |  |
| **TÜ2270** | 56 | 36 | 12 | 20 |  |
|  | 23 | 24 | 11 | 9 |  |
|  | 25 | 90 | 35 | 37 |  |
| sum | 104 | 90 | 35 | 37 |  |
| % | 39.1 | 33.8 | 13.2 | 13.9 |  |
| N=266 |  |  |  |  |  |
| **TÜ2271** | 71 | 2 |  | 1 |  |
|  | 301 | 18 | 3 |  |  |
| sum | 372 | 20 | 3 | 1 |  |
| % | 93.9 | 5.1 | 0.8 | 0.3 |  |
| N=396 |  |  |  |  |  |
| **TÜ2272** | 27 | 7 | 1 | 1 |  |
|  | 295 | 6 | 1 |  |  |
| sum | 322 | 13 | 2 | 1 |  |
| % | 95.3 | 3.8 | 0.6 | 0.3 |  |
| N=338 |  |  |  |  |  |
| **TÜ2273** | 64 | 12 | 2 | 1 |  |
|  | 141 | 39 | 3 | 4 |  |
| sum | 205 | 51 | 5 | 5 |  |
| % | 77.1 | 19.2 | 1.9 | 1.9 |  |
| N=266 |  |  |  |  |  |
| **TÜ2274** | 50 | 22 | 4 | 8 |  |
|  | 141 | 81 | 15 | 16 |  |
| sum | 191 | 103 | 19 | 24 |  |
| % | 56.7 | 30.6 | 5.6 | 7.1 |  |
| N=337 |  |  |  |  |  |

**Table S3:** *albino* k.o.; Cas9 and SpRY proteins

|  | none | weak | moderate | good | very good /<br>complete |
| --- | --- | --- | --- | --- | --- |
| **Cas9** | 1 | 9 | 6 | 15 | 9 |
|  | 1 | 0 | 4 | 4 | 6 |
|  | 5 | 7 | 11 | 11 | 6 |
|  | 2 | 14 | 24 | 35 | 17 |
| **sum** | 9 | 30 | 45 | 65 | 38 |
| **%** | 4.8 | 16.0 | 24.1 | 34.8 | 20.3 |
| N=187 |  |  |  |  |  |
| **aNLS-Cas9** | 0 | 0 | 5 | 6 | 65 |
|  | 3 | 0 |  | 7 | 58 |
|  | 4 | 0 | 1 | 3 | 48 |
| **sum** | 7 | 0 | 6 | 16 | 171 |
| **%** | 3.5 | 0 | 3.0 | 8.0 | 85.5 |
| N=200 |  |  |  |  |  |
| **SpRY** | 0 | 1 | 14 | 20 | 20 |
|  | 0 | 0 | 19 | 16 | 18 |
|  | 1 | 5 | 4 | 16 | 2 |
|  | 5 | 8 | 15 | 58 | 77 |
|  | 43 | 51 | 79 | 97 | 69 |
| **sum** | 49 | 65 | 131 | 207 | 186 |
| **%** | 7.7 | 10.2 | 20.5 | 32.4 | 29.2 |
| N=638 |  |  |  |  |  |
| **aNLS-SpRY** | 4 | 11 | 10 | 23 | 19 |
|  | 0 | 0 | 1 | 8 | 9 |
|  | 4 | 36 | 80 | 85 | 30 |
| **sum** | 8 | 47 | 91 | 116 | 58 |
| **%** | 2.5 | 14.7 | 28.4 | 36.3 | 18.1 |
| N=320 |  |  |  |  |  |

**Table S4:** *albino* repair; SpRY and aNLS-SpRY, OP2* sgRNA, TÜ2270 oligonucleotide donor

|  | none | weak | moderate | good | very good / complete |
| --- | --- | --- | --- | --- | --- |
| **SpRY** | 31 | 17 | 10 | 11 | 0 |
|  | 33 | 20 | 11 | 5 | 0 |
|  | 113 | 41 | 34 | 15 | 0 |
| **sum** | 177 | 78 | 55 | 31 | 0 |
| **%** | 51.9 | 22.9 | 16.1 | 9.1 | 0 |
| N=341 |  |  |  |  |  |
| **aNLS-SpRY** | 21 | 17 | 12 | 20 | 0 |
|  | 47 | 10 | 4 | 10 | 0 |
|  | 95 | 73 | 88 | 56 | 0 |
| **sum** | 163 | 199 | 104 | 86 | 0 |
| **%** | 36.0 | 22.1 | 23.0 | 19.0 | 0 |
| N=453 |  |  |  |  |  |

**Table S5:** *albino* k.o.; aNLS-Cas9 and Cas9 (IDT) 500 ng/μl

|  | none | weak | moderate | good | very good / complete |
| --- | --- | --- | --- | --- | --- |
| **aNLS-Cas9** | 6 | 9 | 23 | 38 | 123 |
|  | 7 | 8 | 28 | 37 | 118 |
|  | 2 | 6 | 8 | 25 | 94 |
|  | 3 | 3 | 7 | 36 | 79 |
|  | 0 | 1 | 4 | 8 | 69 |
| **sum** | 18 | 27 | 70 | 144 | 483 |
| **%** | 2.4 | 3.6 | 9.4 | 19.4 | 65.1 |
| N=742 |  |  |  |  |  |
| **Cas9 (IDT)** | 7 | 6 | 15 | 86 | 131 |
|  | 7 | 8 | 24 | 74 | 131 |
| **sum** | 14 | 14 | 39 | 160 | 262 |
| **%** | 2.9 | 2.9 | 8.0 | 32.7 | 53.6 |
| N=489 |  |  |  |  |  |

